# Quantitative Evaluation of the Cellular Uptake of Nanodiamonds by Monocytes and Macrophages

**DOI:** 10.1101/2022.08.26.505383

**Authors:** Maria Niora, Mathilde Hauge Lerche, Martin Dufva, Kirstine Berg-Sørensen

## Abstract

Nanodiamonds (NDs) with NV^−^ defect centers are great probes for bionanotechnology applications, with potential to act as biomarkers for cell differentiation. To explore this concept, uptake of NDs (~120nm) by THP-1 monocytes and monocyte-derived M0-macrophages is studied. The time course analysis of ND uptake by monocytes confirms differing ND-cell interactions and a positive time-dependence. No effect on cell viability, proliferation and differentiation potential into macrophages is observed, while cells saturated with NDs, unload the NDs completely by 25 cell divisions and subsequently take up a second dose effectively. ND uptake variations by THP-1 cells at early exposure-times indicate differing phagocytic capability. The cell fraction that exhibits relatively enhanced ND uptake is associated to a macrophage phenotype which derives from spontaneous monocyte differentiation. In accordance, chemical-differentiation of the THP-1 cells into M0-macrophages triggers increased and homogeneous ND uptake, depleting the fraction of cells that were non-responsive to NDs. These observations verify that ND uptake allows for distinction between the two cell subtypes based on phagocytic capacity. Overall, NDs demonstrate effective cell labeling of monocytes and macrophages, and are promising candidates for tracking biological processes that involve cell differentiation.

## Introduction

Fluorescent nanoparticles (NPs) are widely applicable tools to answer biological questions, through fluorescence microscopy. They have been used for labeling cellular structures, for treatment of malignant cells and for monitoring biological processes (1, 2). Major challenges when using fluorescent nanoparticles in biological research are their potential cytotoxicity, phototoxicity and photobleaching. Nanodiamonds (NDs) have gained interest due to their high biocompatibility and their uptake mechanism by cells is studied in several works (3–6). In cancer research, NDs have shown great potential as drug carriers and have been successfully loaded with chemotherapeutics adsorbed to their surface, aiming to induce cancer cell death via drug delivery (7–15). So called defect centers convert NDs into photostable and non-photobleachable fluorescent probes (16, 17), making them interesting for cell labeling and time-lapse microscopy (18). Most common are the negatively-charged nitrogen-vacancy (NV^−^) centers which, upon excitation by green light, emit in red and NIR (19, 20). Within the rising field of diamond magnetometry, NDs with NV^−^ defects have been explored as sensors in biological systems to detect temperature variation (21–23), free radicals produced by cells upon stimulation (24–27), and other sensing capabilities (19, 28, 29). These developments demonstrate a potential diagnostic value of NDs with defect centers.

To achieve cellular ingestion of NPs, surface modifications of the particles are often necessary. However, efficient uptake of non-functionalized NDs by various adherent cell cultures, especially HeLa cells, has been reported (4–6, 30–39). Despite their important role in inflammatory diseases and tumor progression, little research has been conducted studying NDs in phagocytic cells (27, 40–42). In this study, we focus on suspension monocytes and monocyte-derived adherent M0-macrophages. Both the un-activated circulating monocytes and un-polarized M0-macrophages are recruited into tissues, before they differentiate into polarized subsets of macrophages (43), often playing a negative key role in disease progression (44, 45). In order to study such processes, the ability of monocytes to differentiate is crucial and should remain unaffected by ND uptake.

Here, we provide a thorough study of the cellular uptake of NDs with NV^−^ defects by monocytes in suspension and their differentiated adherent M0-counterparts, as evaluated by fluorescence microscopy and flow cytometry. First, we time-dependently investigate the interactions of the THP-1 monocytic cell line with unfunctionalized NDs, and quantify their cellular uptake over the course of 3 days. We show that NDs readily interact with the monocytes, without compromising cell viability, proliferation ability and differentiation potential into M0-macrophages. Last, based on the increased ND -uptake and -cellular response by M0-macrophages compared to monocytes, we introduce the notion of using NDs as biomarkers for cell differentiation.

## Results and Discussion

### NDs Uptake by THP-1 monocytic cell line

To evaluate the cellular interactions of monocytes with NDs by fluorescence microscopy, in a time-dependent manner, we subjected THP-1 cells to NDs for up to three days (Fig. 1). In the seeding medium, NDs were 132 ± 62 nm (median ± SD) in diameter, as characterized by nanoparticle tracking analysis (NTA) (Fig.S1). To verify cellular internalization of NDs (red), cells were additionally stained for plasma membrane (green) and nucleus (blue) (Fig. 1A). The middle optical slices display the sub-cellular ND localization for three different exposure-times (left column), and illustrate that ND adherence, internalization and accumulation into monocytic THP-1 cells are time-dependent processes. The related integrated fluorescence signal of the NDs was superimposed on the bright-field images (right column) and verifies the positive correlation of incubation period with ND-cell interactions. Within the first hour, NDs mainly adhered to the plasma membrane, followed by minor local cellular internalization. In the course of time, ND accumulation within the cytoplasm was augmented, while at all time-points, we often noticed additional considerable agglomeration of NDs onto the cell surface. NDs never reached the nucleus, agreeing with literature (40). Notably, some cells exhibited early internalization of NDs dispersed within the cytoplasm. We hypothesized that this was a result of enhanced phagocytic capability. Representative images from all time-points are collectively presented in supplementary Fig. S2.

**Fig. 1.**
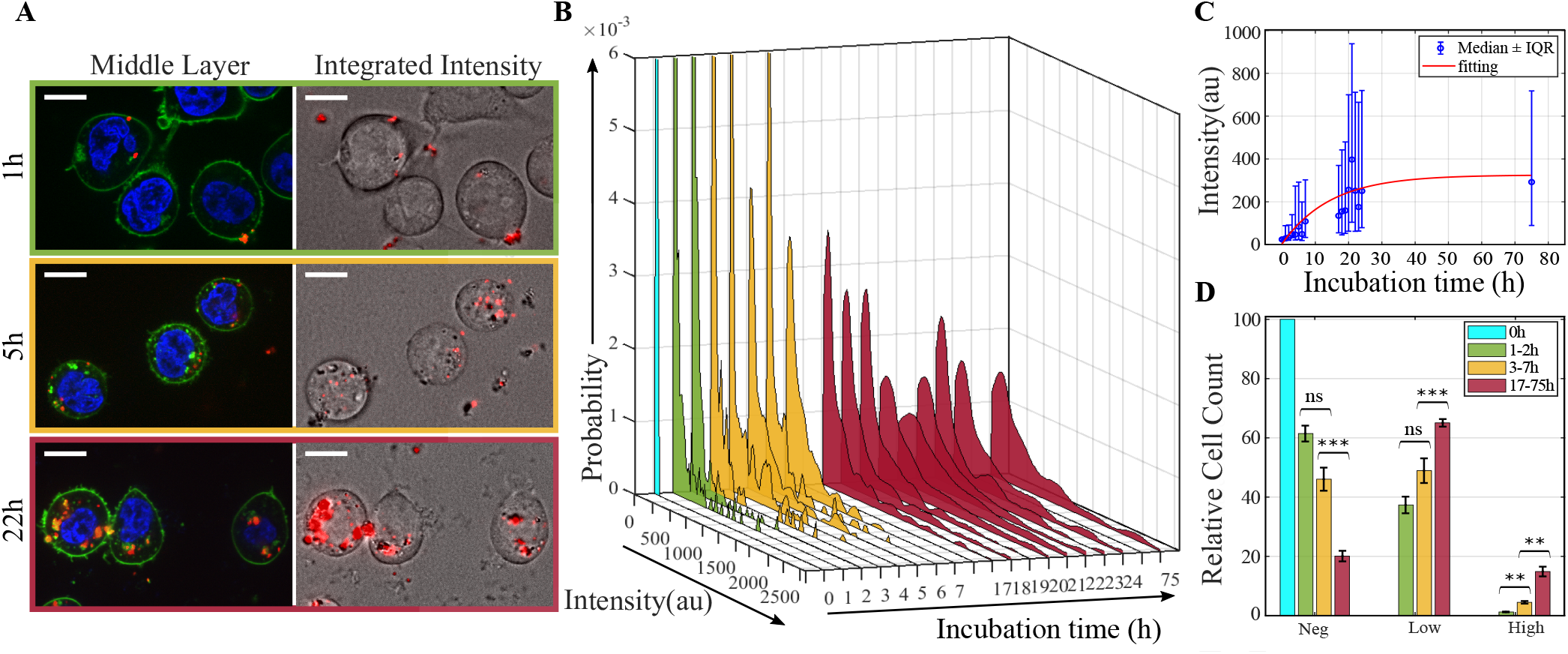
Time-dependent interactions of THP-1 monocytes with NDs (20 μgr/ml). The pseudocolored images of live THP-1 cells treated with NDs (red), and cell stains for nucleus (blue) and plasma membrane (green), were acquired by confocal microscopy (A). The middle optical slices for ND cellular localization (left) and their corresponding integrated ND intensity for cumulative uptake (right), are framed in green, yellow and red for ND administration periods of 1h, 5h and 22h, respectively. The probability densities of the mean ND intensity per single cell against incubation-time (B) were grouped in four time-categories of 0h (cyan), 1-2h (green), 3-7h (yellow) and 17-75h (red). The temporal uptake profile is presented by the medians with IQR, overlaid with exponential fitting in red (C). The differential phagocytic activity of THP-1 cells is clear after dividing the NDs intensity into three ranges, *Neg* < *Low* < *High* (D). Scale bars: 10μm. Statistical significance: Student t-test, α=0.05, **p<0.01, ***p<0.001.

By image analysis, we calculated the mean ND fluorescence intensity per cell, to compute the probability density functions (PDFs), as described in the materials and methods. These PDFs revealed differing interactions of THP-1 cells with NDs and confirmed a positive temporal-dependence of ND uptake 1B). NDs were administered to the cells for different exposure-periods, spanning from 1h up to 75h. At all time-points, we monitored a wide range of ND intensity including both a cell population with mean ND intensity similar to controls (0h of NDs), which indicates lack of ND-cell interactions, but we also detected a cell population with increased ND fluorescence, suggesting enhanced cellular uptake. Uptake variations, especially at the earlier time-points, suggest variation in phagocytic activity or could be a result of ND size dispersity. Next, the temporal uptake profile, obtained by plotting the medians with interquantile ranges (IQR) (Fig. 1C)), showed that the ND-monocyte interactions saturate exponentially with incubation-time and seemingly approach a plateau at around 14h. This is most likely a result of saturated cellular uptake capacity, in combination with both (i) cellular ND excretion and cell proliferation, leading to fluorescence dilution, and (ii) the ongoing exposure to NDs, which are competing processes.

We further grouped the distribution profiles into four time-categories (0h, 1-2h, 3-7h and 17-75h), as highlighted with cyan, green, yellow and red (Fig. 1B). For each of them, we integrated the histogram absolute frequency counts, in three main ranges of ND intensities, denoted as *Neg*, *Low* and *High*, as detailed in the materials and methods. We then calculated the relative frequency counts per ND-range, for each time-group (Fig. 1D). This analysis demonstrated that the THP-1 cells take up the NDs readily (37% in *Low*, green), with the fraction of cells in the *Neg* range decreasing over time, while a respective increase is observed by cells of *Low* (65% in *Low*, red) and *High* fluorescence. Interestingly, even after 75h of uninterrupted exposure to NDs, cells in the *Neg* range were not eliminated (12%), which positively contributes to the saturation of the temporal uptake profile. A positive temporal dependence of ND interactions with THP-1 monocytic cells and a cumulative ND uptake with exponential saturation corroborates similar observations with silver NPs in THP-1 cells (46) and with NDs in primary human monocytes (40).

The amount of cellular particle internalization depends on particle properties, which affect the mechanisms of particle uptake. Monocytes are expected to both phagocytose and endocytose particles, as demonstrated by Gatto et al. with 5nm NPs in THP-1 cells, using TEM microscopy (47). Due to the large size of the NDs we used, clathrin-mediated endocytosis is a likely mechanism to take place (48), as reported previously with similar-sized NDs administered to non-phagocytic cell lines (3–6) and to phagocytic macrophages (41, 42). Contrary, caveolae-mediated endocytosis has been suggested to occur with NPs smaller than 80nm (49), hence this process is unlikely herein.

### Uptake Saturation and ND Biocompatibility

To test if cells could be saturated by NDs, we exposed THP-1 monocytes to varying concentrations of NDs, for 24h (Fig.2). The fluorescence intensity of NDs was evaluated by single cell analysis, on data sets from two different experimental methods. The results from both optical microscopy (Fig. 2A) and flow cytometry (Fig. 2B) showed that ND uptake was not dose-dependent, for the tested concentrations, and it appears we reached an uptake capacity limit. The ND distribution profiles from flow cytometry were verified to be comparable to those obtained by optical microscopy (Fig. S4B).

**Fig. 2.**
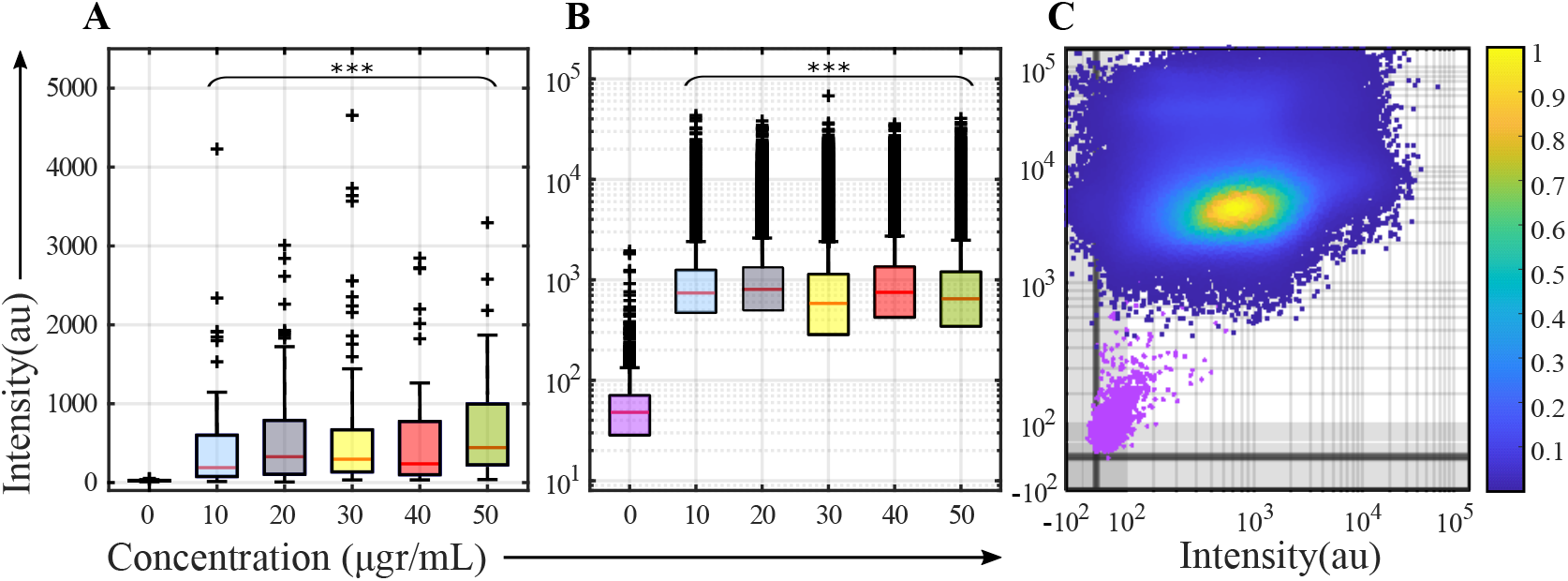
Response of THP-1 monocytes to different concentrations of NDs for 24h. ND uptake was determined by single-cell analysis of ND fluorescence. For optical microscopy, we report the mean values (A) and for flow cytometry, we report the integrated values (B). The intensity of NDs against viability stain was evaluated by flow cytometry (C). Results are compared to untreated controls (purple). Statistical significance: Kruskal-Wallis test, α=0.05, ***p<0.001.

Our findings are not consistent to reports on ND dose-dependence studies in adherent non-phagocytic cell lines (32, 38), adherent macrophages (41) and suspension primary human monocytes (40). Discrepancies with other monocyte investigations, may result from the difference in monocytic cell origin, since the immortalized cell line that we used shows heterogeneous uptake populations. Another important difference derives from the different incubation protocols. In contrast to Suarez-Kelley et al., we used round bottomed wells that collect both the cells and the NDs in a small area, likely increasing uptake efficacy. In their study, the flat bottomed wells (common in adherent cell cultures) disperse the cells onto the surface, hence higher concentrations are required to achieve similar final NDs-to-cell ratio. Another underlying reason is the different size of NDs that lead to different mechanisms of cellular uptake into play. In the Suarez-Kelly study, NDs were around 70nm, a particle size which is known to allow for caveolae-mediated endocytosis (46) whereas the larger NDs we use allow for clathrin-mediated endocytosis and further ND agglomeration during culture facilitates phagocytosis. In their case, potentially by increasing the ND concentration, ND size increases by agglomeration triggering additional uptake mechanisms. The above suggest that the culture protocol and ND uptake mechanism could influence the dose-dependence of ND cellular uptake.

ND biocompatibility was investigated by additional cell staining with a cell-permeant viability stain. We did not observe a cytotoxic effect despite the increasing ND exposure-dose, as demonstrated by the scatter plot of ND fluorescence intensity vs viability stain, per single-cell, obtained by flow cytometry (Figure 2C). All cells were positively stained with the viability stain, in contrast to the unstained control (purple), while the distributions of viability stain were identical for all ND doses (Figure S4A). That ND biocompatibility is dose-independent is also demonstrated in other similar dose-dependent ND-cytotoxicity studies (34, 38, 40–42, 50). From the microscopy data, we verified a positive linear correlation between the ND uptake and cellular viability, by Spearman correlation coefficient (ρ = 0.78) (details can be found in the Supplementary Figure S4C). This implies that ND cellular uptake reflects phagocytic activity of the cells, taking cell viability as an indication of general cellular activity.

### ND Uptake by M0-macrophages

We investigated the ability of macrophages to ingest NDs, after chemically-differentiating the monocytic cell line into macrophages by PMA. Using flow cytometry immunophenotyping analysis, we verified the cells’ altered phenotype towards the non-polarized M0-subtype of macrophages. This was evaluated by the expression of cell surface markers CD13 and CD14, after binding of respective monoclonar antibodies (see Supplementary Fig. S5).

To evaluate the cells uptake capability, we employed confocal microscopy and flow cytometry. Confocal images of macrophages exposed to NDs (red) for 6h, and co-stained for nuclei (blue) and cell membranes (green) revealed that NDs distributed broadly and seemingly more uniformly inside the cytoplasm (Fig. 3A), compared to most THP-1 cells. The integrated ND fluorescence signal (Fig. 3B) verified that NDs interact with the macrophages to a greater extent than with the monocytes (comparison with Supplementary Fig. S2). In regards to cell morphology, the macrophages exhibited increased cytoplasmic to nuclear ratio compared to the non-differentiated THP-1 cells.

**Fig. 3.**
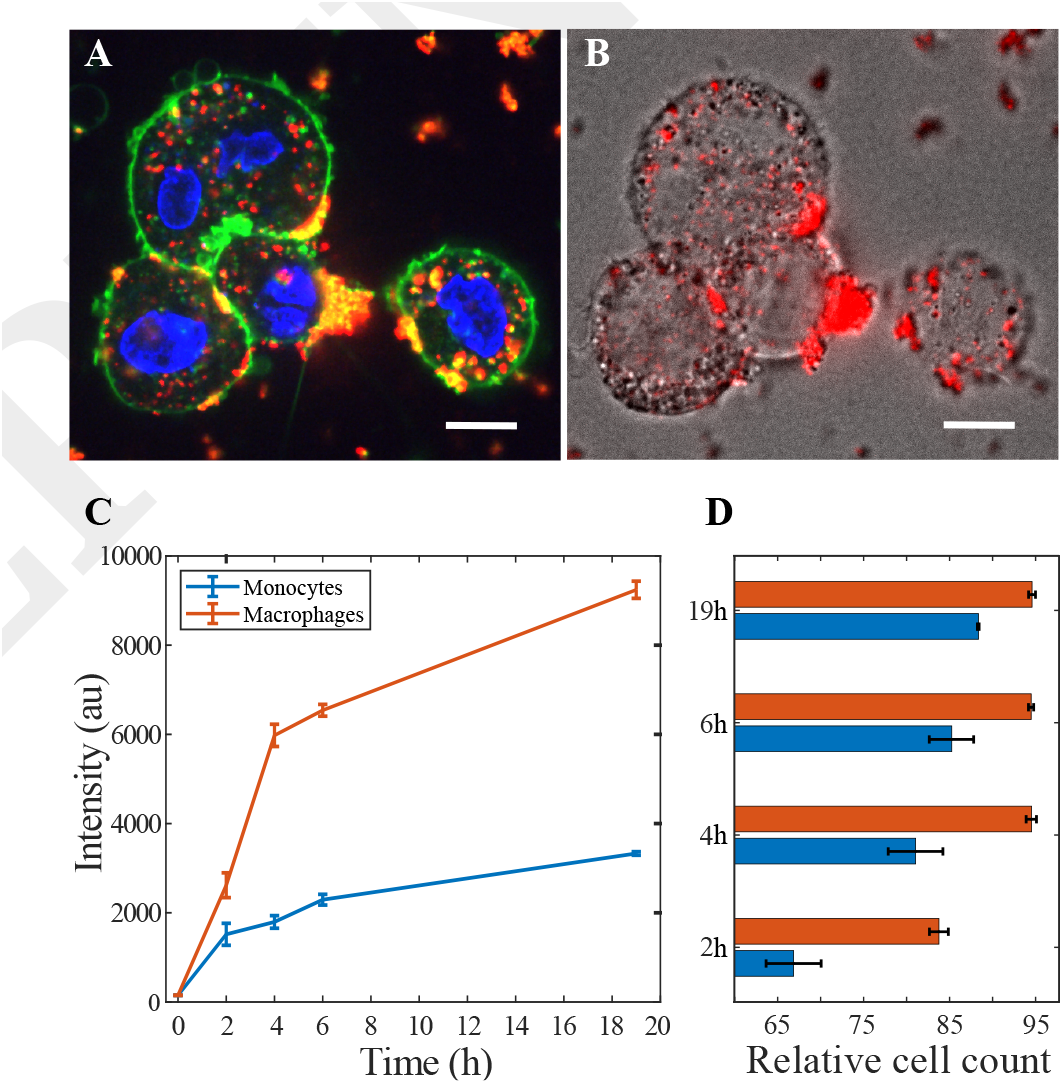
ND uptake by chemically-differentiated M0-macrophages. Fluorescence microscopy of cells labeled with NDs (red) for 6h (20 μgr/ml), plasma membrane stain (green) and nuclear stain (blue) (A), and the overlay of the integrated ND fluorescence with bright-field (B). Temporal ND fluorescence by M0-macrophages (red; 745,000 cells) and THP-1 cells (blue; 1,400,000 cells), as assessed by flow cytometry (median ± SEM, n=4) (C). The percentage of cells with positive response to NDs over time (median ± SEM) was greater for the macrophages (D). Scale bars: 10μm.

For quantitative comparisons, we assessed cellular ND uptake by flow cytometry after ND administration for different incubation-periods. Both the THP-1 monocyte-like cells (red) and the monocyte-derived macrophages (blue) take up NDs efficiently and increasingly with time (Fig.3C). The macrophages displayed comparably significant enhanced ND uptake, in agreement with similar works employing different NPs (51). Temporal increase in macrophage uptake is presented also by others (41, 42) with NDs of similar size. Our evaluation was based on frequency histograms of integrated ND fluorescence intensity per single cell (Fig. S6), from which we calculated a median value for each time-point. The percentages of NDs(+) cells were greater for the macrophages (red) over the THP-1 monocytic cell line (blue) at all time-points (Fig.3D), verifying enhanced macrophage response to the NDs. To gate NDs(+), the threshold was determined from untreated controls (0h of NDs).

The cell populations that phagocytose the most, were detected by flow cytometry analysis of light scattering on cells without (A, B, C) and with (D, E, F) PMA treatment, before (A, D) and after (B, C, E, F) administration of NDs (Fig.4). In agreement with the microscopy images, PMA-treatment renders the macrophages greater in both size and granularity, as a result of the altered phenotype. Consequently, it increases the fraction of cells with high FSC-A and SSC-A, while being double-positive for CD13 and positive for CD14 (the population outside the blue dashed gates). The small fraction of such cells that is detected in the control THP-1 cells (no PMA) reveals spontaneous differentiation into macrophages during normal culture conditions. The gating strategy of the monocyte-like populations is explained in Supplementary Fig. S4. THP-1 monocytes are reportedly prone to spontaneous cell differentiation into a more mature phenotype during culture (52). We speculate that this variation in phenotype is associated to phagocytic activity, and would explain the increased dispersion of ND uptake, at the early time points (Fig.1). Cells with macrophage phenotype took up larger amounts of NDs (Fig.3C), while increasing the fraction of macrophages resulted in a respective increase in the number of cells with enhanced ND uptake capacity (Fig. 4; B/C versus E/F) A larger fraction of monocytes (no PMA) remained NDs(−) (23% for monocytes over 5% for macrophages). We thus assume that macrophages exhibit higher phagocytic activity compared to monocytes, as PMA-treatment depletes the population of NDs(−). The histograms of ND fluorescence in the Supplementary Fig. S6) show clearly this population depletion, which is more profound at the early exposure-points, supporting our presumption. ND phagocytosis by macrophages is confirmed elsewhere with phagocytosis inhibitors (41) and TEM microscopy (53).

**Fig. 4.**
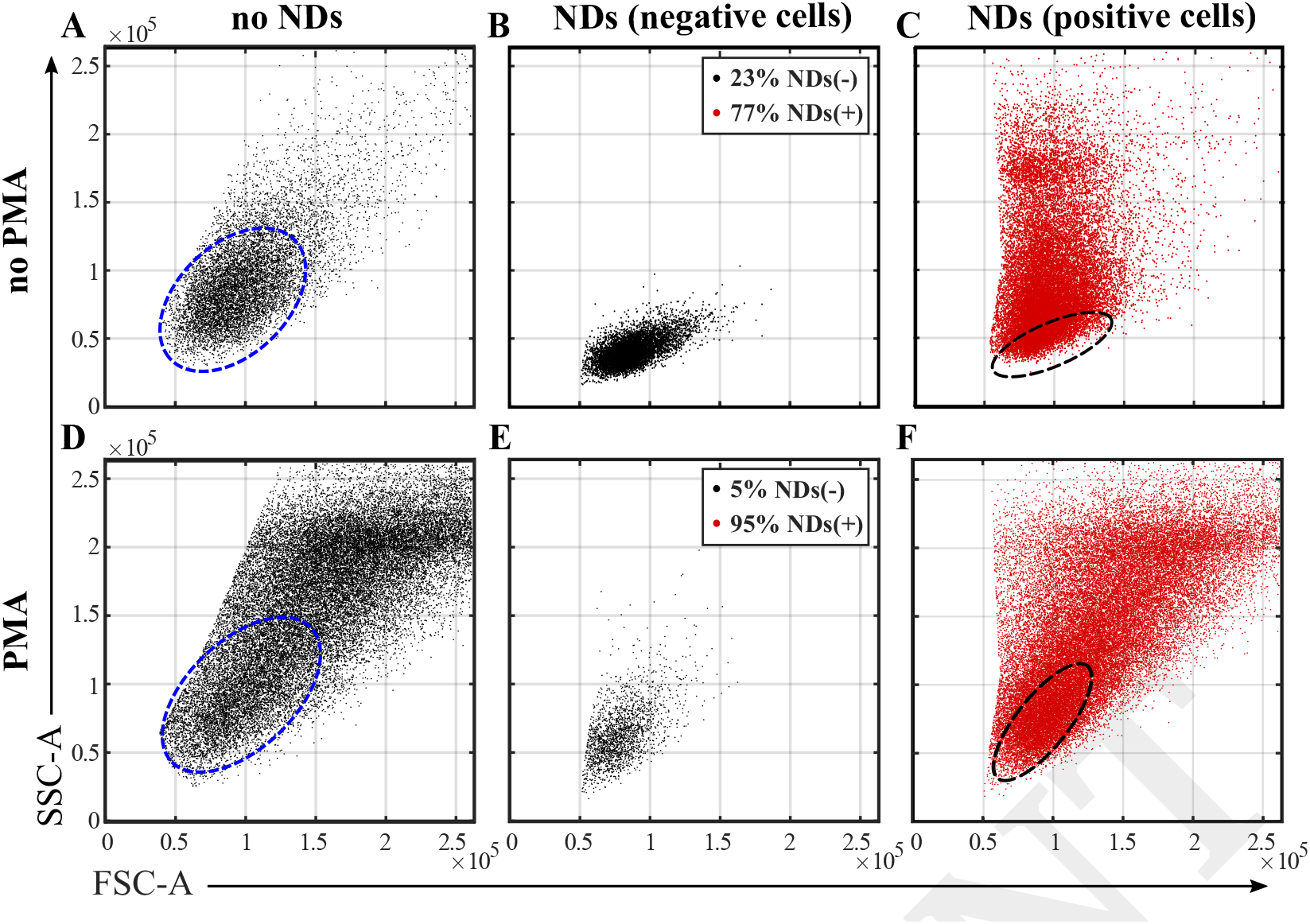
Flow cytometry analysis of THP-1 cells prior to- (A,B,C) and post- (D,E,F) PMA-induced cell differentiation. Cells were examined both before- (A,D) and after- exposure to NDs for 4h. Before exposure to NDs, we identified the monocytic population by antibody staining by CD13 and CD14 (blue dashed ellipses), versus the macrophage population (outside the gates). After ND administration, we distinguished between the cell populations with a positive or negative response to NDs, marked in black for NDs(−) and in red for NDs(+), respectively. The black dashed ellipses highlight the area of NDs(−).

Due to protocol practicalities, we did not perform both antibody labeling and ND-administration in the same samples. However, the above gating strategy allows us to indirectly detect macrophage populations in non-labeled samples. Notably, in response to ND uptake, the monocytes, which are normally agranulocytes, gain granularity expressed by an increase in side-scatter, incommoding the detection of macrophage population (Fig. 4). By checking cell autofluorescence (upon excitation at 405nm), we verified an increase post-differentiation, as also reported elsewhere for cells subjected to PMA-treatment (54). We observed a similar pattern in THP-1 cells that were spontaneously differentiated without external stimuli. The cells of higher size (FSC-A) and granularity (SSC-A), additionally expressed both enhanced response to NDs and cell autofluorescence (data not shown), supporting our suggestion for phenotype-distinction without antibody labeling.

### Effect of ND Uptake on cellular functions

We were interested to know whether ND uptake affects major cellular functions. We first investigated a potential impact on cell proliferation. We maintained two parallel cultures of THP-1 cells that were “ND-labeled” by 24h exposure (i) and “non-labeled” (ii), and subsequently subcultured as normal. We measured similar cell density and viability for both conditions, at every passage over a period of three weeks, with no evidence of cytotoxic effect or cell growth inhibition induced by the ND uptake. Similar to our observations, Thomas et al., who investigated ND impact on macrophages’ proliferation, in a size-dependent manner, reported no effect for NDs larger than 100nm (53), while others demonstrated no impact on cell division for various kinds of cells by ~100 nm NDs (6, 36). After 5 subcultures (25 cell divisions), we examined both (i) and (ii) by flow cytometry, and evaluated the ND fluorescence signal (Fig. 5A). At this point in culture, the cells had lost the NDs, as illustrated by the identical zero ND-expression for both conditions (blue curves). It has been previously demonstrated, with several non-phagocytic cell lines, that during cell mitosis the daughter cells share the internalized NDs of the parent cell, until no NDs are observed in the cells several generations later (6, 33). To assess whether this process affects cellular uptake capability, we re-exposed the cells to NDs for 2h. We observed similar ND uptake (red curves) by both the cells exposed to NDs from condition (ii), and the cells from condition (i) that lost the initial NDs and were now re-exposed to NDs. This is indicative that NDs-treatment does not impair long-term cellular uptake capability.

**Fig. 5.**
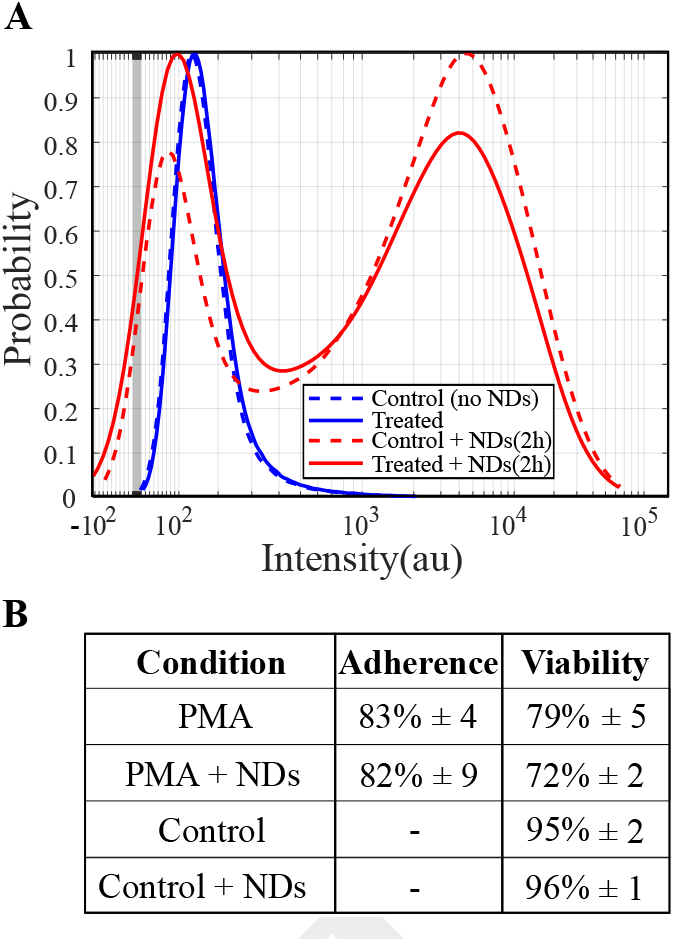
Effect of ND Uptake on cellular functions, i.e., cellular uptake (A) and cell differentiation (B). ND fluorescence in single cells that were exposed to NDs for 24h followed by 25 cell divisions (solid blue curve) reveals that NDs efficiently exited the cells. Subsequent re-exposure to NDs for 2h (solid red curve) verifies the unaffected long-term uptake ability, as compared to controls (dashed curves). Cell characteristics (averages ± SEM), for both PMA-treated (n=3) and controls (no PMA), show that cells’ pre-exposure to NDs for 24h (+ NDs) did not affect the differentiation potential.

We last investigated whether ND uptake had an impact on cell differentiation, by repeating the same 5-days differentiation-protocol to THP-1 cells post-0h (PMA) or 24h (PMA + NDs) of -exposure to NDs (Fig. 5B). After 3-days culturing in PMA, non-adherent cells were subjected to hemocytometer analysis, upon which we verified similar level of cell adherence. Two days later, when PMA-treatment was completed, we harvested and evaluated the adherent cells. Again, the similar cell densities and viabilities, indicated no restriction in cell differentiation ability. The morphology of the cells was also confirmed to effectively change into adherent culture, by inspection with an inverted light microscope. Our findings agree with another study using the same differentiation protocol but exposing the THP-1 cells to a different type of NPs (47), and with a study using NDs during cell differentiation of primary murine bone marrow cells into macrophages (41). Cell viability after PMA-treatment appears lowered throughout the protocol, due to inhibition of cell proliferation induced by the chemical. The expected arrest in proliferation (51), was verified by cell density counts at the end of PMA-treatment, with a 7-fold decrease relative to controls (no PMA). Based on cell adherence, cell viability and proliferation inhibition we concluded that the cell differentiation potential remained unaffected.

## Conclusion

THP-1 monocytes are a good cell model to study immune cell interactions with their environment, thanks to their ability to differentiate into more mature phenotypes. In order to study their activity we have employed carboxylated NDs with defect NV^−^ centers, and we investigated ND uptake by both monocyte-like THP-1 cells and their actively-differentiated unpolarized counterparts; M0-macrophages. The cells capability to engulf NDs of 132 ± 62 nm in diameter was based on fluorescence emitted from the NV^−^ centers and was evaluated by single cell analysis using confocal microscopy and flow cytometry.

We studied ND uptake by monocytes in a time-dependent manner and we found differing ND-cell interactions, with a temporal dependence, which is attributed to ND agglomeration onto the plasma membrane and accumulation within the cytoplasm. The notable varying level of ND internalization at early time-points was attributed to uneven phagocytic capability of the cells. We showed that this variation in uptake capacity results from heterogeneous monocyte populations, due to spontaneous cell differentiation without external stimuli, prior to ND exposure. To support our claim, we compared ND interactions with THP-1 monocyte-like cell line to M0-macrophages obtained by PMA-differentiation of the THP-1 cells. Overall, both phenotypes exhibited efficient and increasing temporal ND uptake, but the increased phagocytic activity of the macrophages enhanced their capability to internalize the NDs, and decreased the heterogeneity on ND uptake, which was observed in the case of THP-1 cells. Our findings suggest that the spontaneous THP-1 cell differentiation in culture positively affected ND internalization. We thus reveal a promising usage of NDs to act as markers for cell differentiation.

Last, we demonstrated that ND uptake and biocompatibility were dose-independent, while uptake did not hinder major cell functions, i.e., proliferation and differentiation ability. THP-1 cells were able to totally get rid of the NDs upon normal cell culture for several cell divisions, without affecting their viability, proliferation rate and ability to uptake a second round of NDs. Consequently, we highlight that the studied NDs can safely be utilized as probes for cell labeling and long-term tracking of biological processes, which combined with ND quantum sensing, makes them promising candidates for theranostics, particularly for therapies targeting monocytes and macrophages.

## Materials and Methods

### Cell Culture

THP-1 human monocytic cell line was purchased from American Type Culture Collection (ATCC; TIB-202) and was grown at 37° C, 5% CO_2_, 100% humidity. Cells were cultured in Roswell Park Memorial Institute (RPMI) 1640 medium modified with 2 mM L-glutamine, 10 mM HEPES, 1 mM sodium pyruvate, 4500 mg/L glucose, and 1500 mg/L sodium bicarbonate (Fisher Scientific; ATCC modification), and supplemented with 10% fetal bovine serum (FBS), 100 U/mL penicillin and 100 μg/mL streptomycin (P/S) (Biowest).

### Cellular Uptake of Nanodiamonds

Surface-carboxylated nanodiamonds with 3 ppm nitrogen-vacancies, corresponding to 1200NV^−^ per particle, were 120nm in diameter (Sigma-Aldrich, 798088; Exc/Em: ~560/~680 nm).). Our protocol for ND administration was optimized based on control experiments, varying culture wells, cell density, culture medium and ND treatment pre-administration. THP-1 cells were suspended in fresh seeding medium (FluoroBrite; Gibco, supplemented with 2 mM L-glutamine, 10% FBS and 1% P/S), and were seeded in round-bottom 96-well plates, at a volume of 100μL/well and a concentration of 500,000 cells/mL. NDs were subjected to bath-sonication for 10 min before administration to the cells (20 μgr/mL), for incubation times ranging from 1h to 75h. To investigate cellular uptake capacity we additionally tested different concentrations of NDs (10-50 μgr/mL). Cells were further treated with Cellmask green plasma membrane stain (Invitrogen; Exc/Em: 522/535 nm) and Hoechst 33258 nuclear stain (AnaSpec Inc; Exc/Em: 352⁄461 nm). NDs uptake and localization were evaluated via live-cell imaging by spinning-disk (SD) confocal-microscopy (Nikon Ti2) with a 100× oil objective (CFI Plan Apochromat Lambda, NA 1.45, WD 0.13 mm), followed by image analysis. To study NDs biocompatibility, cell viability was assessed with cell-permeable viability stain calcein UltraBlue AM (Cayman Chemical, Exc/Em: 360/445nm), by both SD confocal-microscopy and flow cytometry (BD Biosciences; LSRFortessa™). For both techniques, final samples were suspended in fresh Fluorobrite-based seeding medium, which effectively enhances fluorescence signal (55), without compromising cell viability, during live-cell imaging. For flow cytometry, cells were filtered through Falcon^®^ cell-strainers of 35 μm mesh (Corning), before examination.

### Cell Differentiation

For cell differentiation into M0-macrophages, THP-1 cells were incubated (37° C, 5% CO_2_, 100% humidity) in 6-well plates with 20 ng/mL phorbol 12-myristate 13-acetate (PMA; Sigma-Aldrich) at a volume of 2mL/well and cell density of 250,000 cells/mL, for 3 days, followed by 2-days rest in fresh culture medium. At the end of the treatment, adherent cells were either labeled with antibodies or exposed to NDs. For immunophenotyping analysis by flow cytometry, cells were detached with dissociating enzyme TrypLe Express (Gibco) and cell scrapers, before applying mouse anti-human monoclonal antibodies against CD13 (eBioscience; Exc/Em: 488/710 nm) and CD14 (eBioscience; Exc/Em: 405/436 nm). To assess cellular uptake, differentiated cells were washed with PBS, exposed to NDs in fresh seeding medium, before further PBS washing and harvesting by cell scraping. Images were collected by SD confocal-microscopy, and for quantitative analysis we used flow cytometry. Control samples were examined for cell viability with either calcein UltraBlue AM or calcein AM (Invitrogen; Exc/Em: 360/445nm).

### Image Pre-processing

For image pre-processing, we used open-source software Fiji (56), for controlled cell selection prior to image analysis. We worked with 50 optical-slices per z-stack, corresponding to 10μm, starting from the cells’ bottom layer upwards, as most monocytes sediment after about 30min rest. For each z-stack, we created a 2D image of z-integrated intensity of NDs, and a 2D binary image based on either plasma membrane stain or viability stain, restraining the analysis to cell surface. That being said, ND agglomerates onto cell surface were only partially included in the subsequent average intensity calculation. A representative example of final blobs’ boundaries is illustrated in Supplementary Fig. S4C. From the binary images, we excluded all unhealthy cells. To identify cells with compromised viability, we visually inspected the bright-field channel for cell boiling-surface, and the fluorescence channels for disturbed membrane staining and/or abnormal nuclear staining (saturated or dotted-pattern). For the ND biocompatibility analysis, we skipped cells’ health state pre-selection.

### Image Analysis

For image analysis we used custom Matlab scripts (MATLAB R2020b; MathWorks). We performed two analyses; one for ND uptake calculation and one for correlation of ND uptake to cell viability (see Supplementary Fig. S4C.). Based on the binary images, we calculated the mean ND fluorescence signal in single-cells and the mean fluorescence intensity of the viability stain, when applied. In both cases, we restrained the analysis to the cell surface, including ND adsorption to the cellular membrane but excluding ND agglomeration beyond the membranes. From the analyses, we excluded cells that were partly outside the field-of view (image borders).

### Probability Density Functions

Based on the mean ND fluorescence intensity per cell, we obtained the probability distributions from more than 90 cells and up to five independent experiments per time-point. From these frequency distributions, we computed the continuous probability density functions (PDFs) by non-parametric kernel density estimation, which reveal the differential interactions of NDs with single live THP-1 cells, in the course of time. At earlier time-points, the mean intensities follow an inversed Gaussian distribution profile, whereas after 17h, the mean intensities follow more of a poisson-like profile, as obtained by probability distribution fitting. That confirms that numerically large values are more probable with increasing incubation. The probability distributions diverge increasingly from normality and they are heavily right skewed, with increasing dispersion. PDF characteristics are summarized in Supplementary Fig. S3. ND fluorescence was further split into three ranges based on PDF characteristics. The three ranges describe the ND-cell interactions, as ‘negative’, *N eg*, based on the control (0h of NDs), as ‘low’, *Low*, spanning above the control up to a cut-off value of 1000, and as ‘high’, *High*, ranging from 1000 up to max value of 23000, all in arbitrary units.

### Flow Cytometry Data Analysis

Flow cytometry data was processed with Matlab software (MATLAB R2020b; MathWorks). Data was loaded with the FCS data reader package (57), followed by custom scripts for gating and numerical analysis. When relevant, the logicle transformation package (58) was also employed.

## Acknowledgements

This work was supported by Independent Research Fund Denmark (grant no 0135-00142B) and the Novo Nordisk Foundation (grant no NNF20OC0061673). MN acknowledges Bente Rotbøl for technical support on flow cytometry.

## Author Contributions

MN conceived the study under supervision of MD, MHL, and KBS. MN conducted all the experiments, prepared the figures and drafted the manuscript. All authors edited and revised the manuscript.

## Data Availability Statement

The data that support the findings of this study are available from the corresponding author upon reasonable request.

