## Supplementary Information for "Quantitative Evaluation of the Cellular Uptake of Nanodiamonds by Monocytes and Macrophages"

**A. NDs in THP1-monocyte-like cell line**

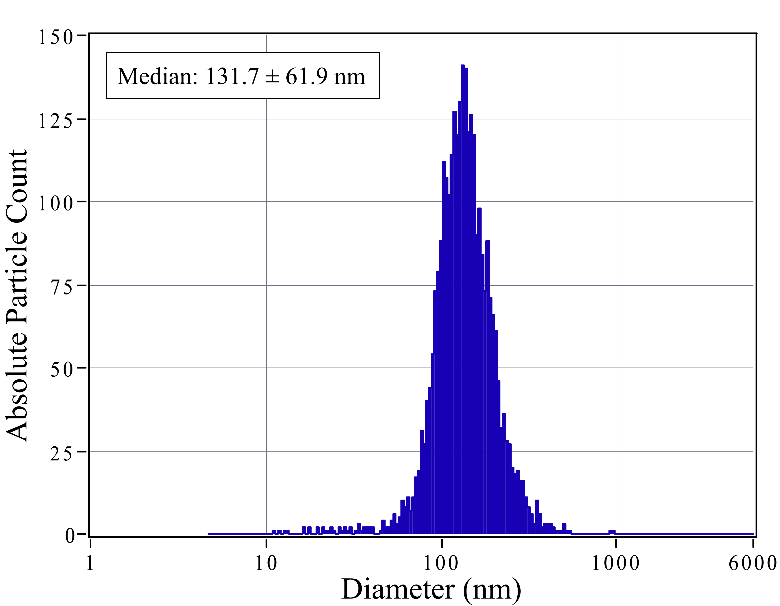

**Figure S1.** Size distribution of the NDs by NTA. NDs (median ± SD, n=11) were sonicated for 10min before suspending into the seeding medium.

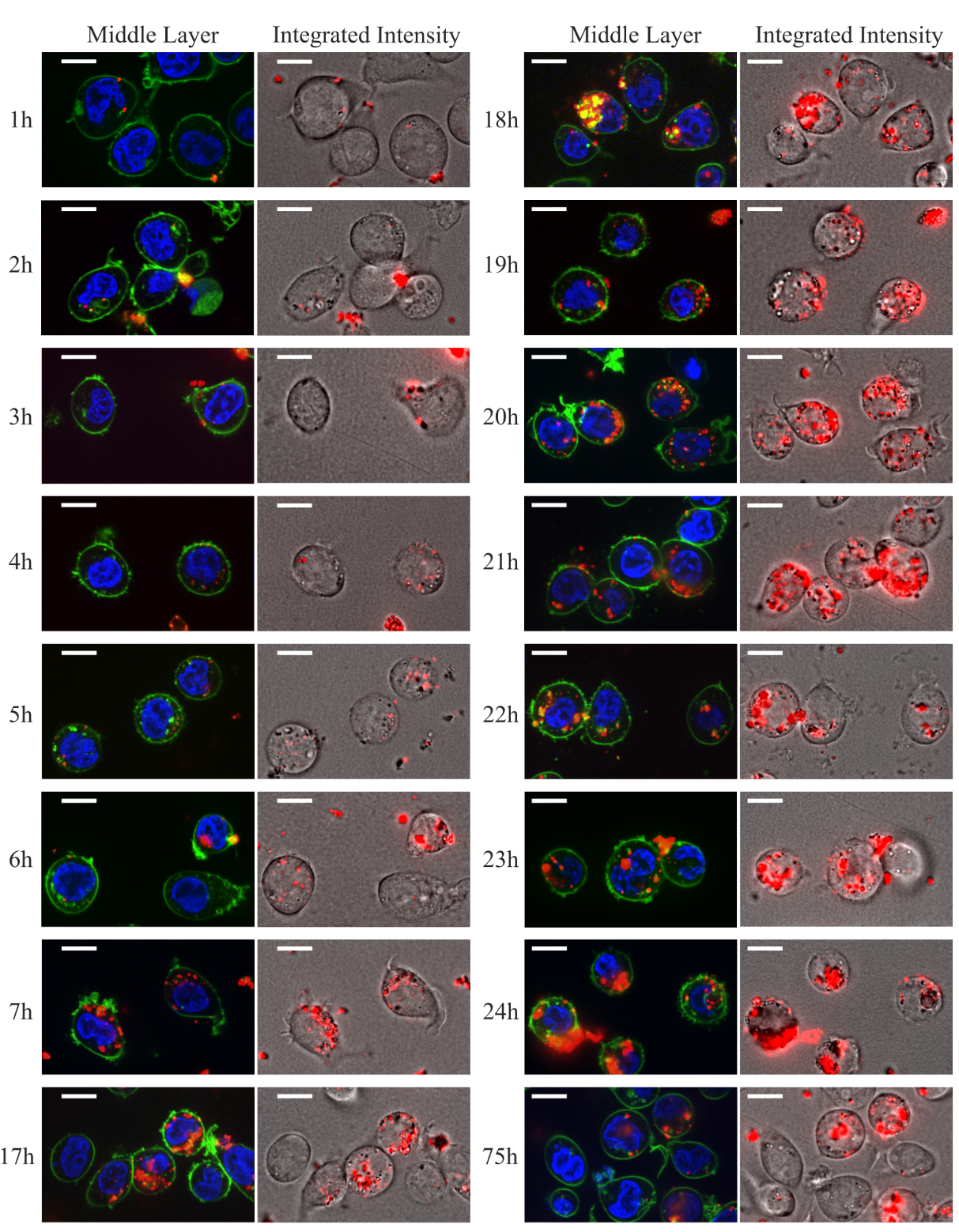

**Figure S2.** Pseudocolored confocal-microscopy images of live THP-1 cells exposed to NDs (red) for up to 75h, and additionally treated with cell stains for nucleus (blue) and plasma membrane (green). For each exposure-period, a middle optical slice (left) is paired with the corresponding integrated NDs intensity (right), to display NDs' subcellular localization and the cumulative ND uptake, respectively. The cellular interactions with the NDs and NDs’ cellular internalization show a positive time-dependence. Scale bars: 10μm.

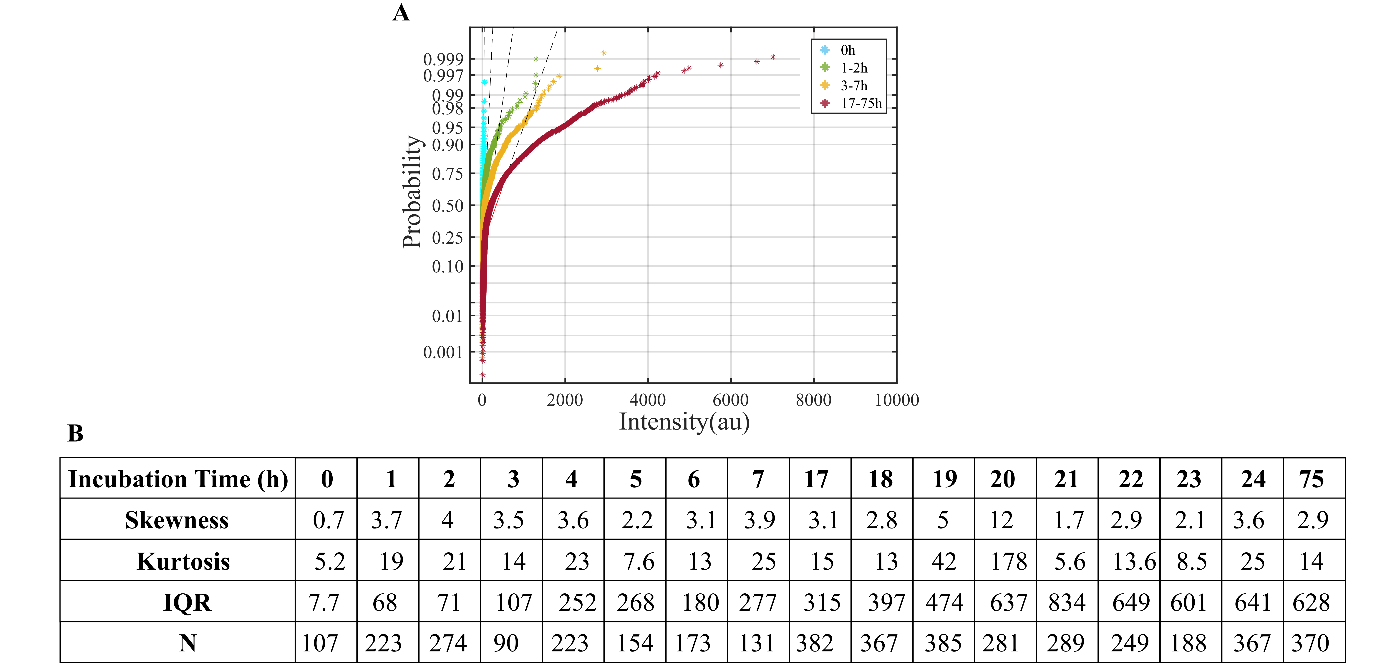

**Figure S3.** Characterization of the probability densities of the mean ND intensity per single THP-1 cell, in the course of time. The normal probability plots show that the density distributions diverge from normality over time, with an increasing right tail (A). The table displays characteristic properties for each density distribution, i.e., skewness, kurtosis and interquantile range (IQR), from a corresponding number (N) of analyzed cells (B).

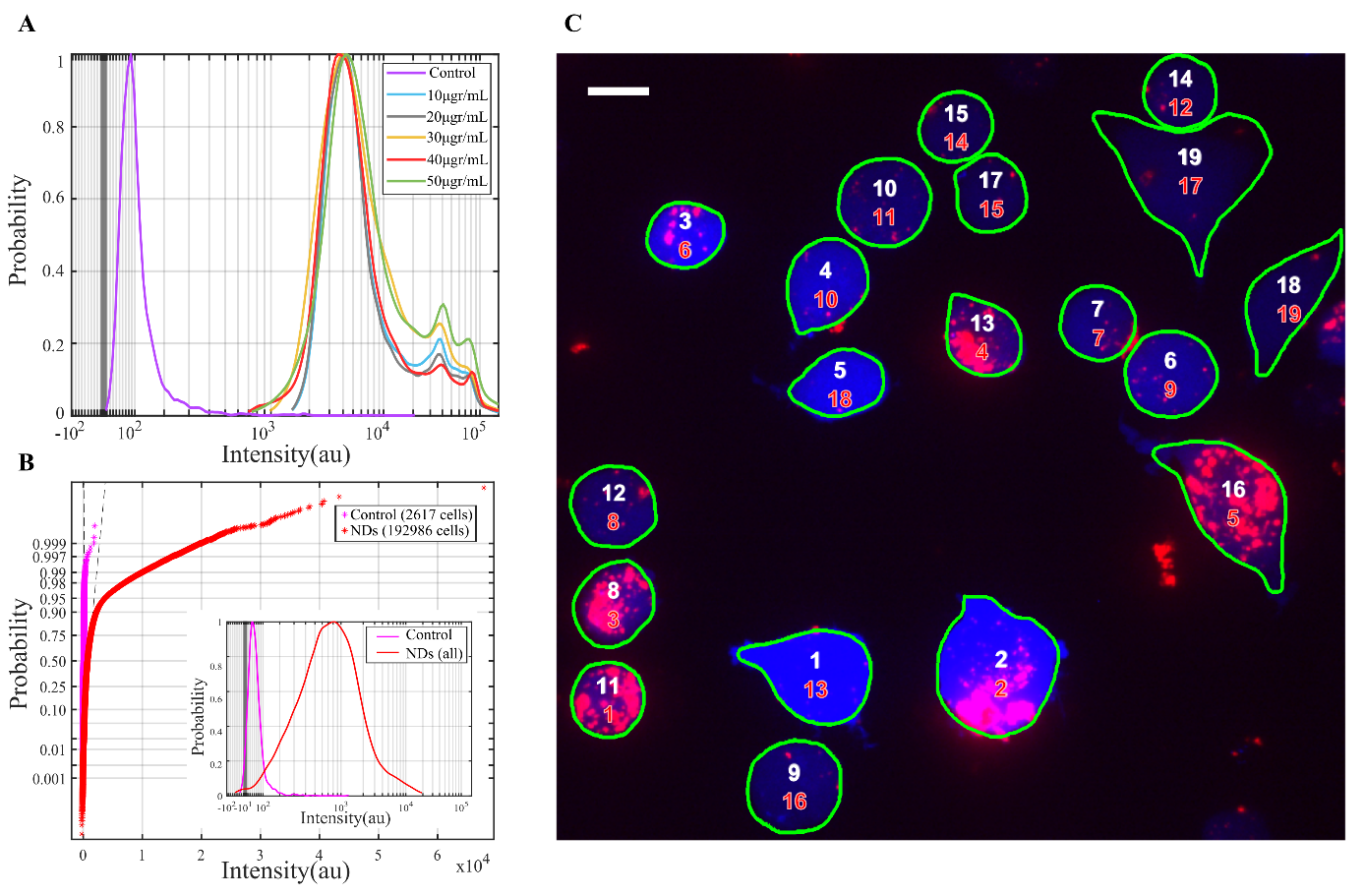

**Figure S4.** Characterization of ND uptake against cell viability, by flow cytometry and confocal microscopy. Cells were exposed to different concentrations of NDs, for 24h, and were additionally labeled for viability by calcein UB AM. Flow cytometry analysis of the viability stain fluorescence (A) shows positive staining for all cells as compared to the unstained control (purple) and identical distribution profiles regardless of the administered concentration of NDs. The normal probability plots (B) of the distribution of ND fluorescence (inset) reveal divergence from normality, with a right tail. Image analysis (C) of cells with NDs (20μgr/mL) by sequential numbering of increasing fluorescence signal for the NDs (red) and the viability stains (white). Single cell analysis was based on detection of the blob borders (green), which point out the boundaries of the cell membranes. Scale bar: 10μm.

**B.** **PMA-induced differentiation**

We actively differentiated monocytes into macrophages to compare their ability to ingest the NDs. THP-1 cells were differentiated into M0-macrophages by 3-days treatment with PMA, followed by 2-days rest in fresh culture medium. Post-treatment, adherent cells were harvested and labelled with antibodies CD13 and CD14, for immunophenotyping with flow cytometry **(Figure S5)**. Compared to untreated controls (no PMA), PMA treatment (PMA) resulted in cells of increasing granularity and size, as observed by the side- and forward- scatter, respectively **(Figure S5A)**. To identify M0-macrophages by CD13 and CD14 we evaluated the fluorescence signals. Monocytes and macrophages co-express the antigens, but expression is higher for macrophages reflecting an increase in cell maturity. Despite the long-term treatment and the overall increased antigen expression post-PMA, there is a remaining population that retains similar expression levels to the controls **(Figure S5; B, C)**. Likewise for the PMA-untreated antibody-stained controls, there is small fraction of cells with size-to-granularity ratio and CD14 expression similar to the M0-macrophages. Gating allowed us to identify the cells that are negative for CD13 and CD14 (black dashed vertical gates), as compared to unstained controls. Then, by backgating the negative-antibody populations on the scatter plots (black dashed elliptic gates), we obtain the size-to-granularity ratio of the monocytes vs differentiated macrophages. We see that all monocytes are at a restrained range, but not all cells in that range were negatively-stained with antibodies, denoting that, despite chemical treatment, some cells preserve their morphology.

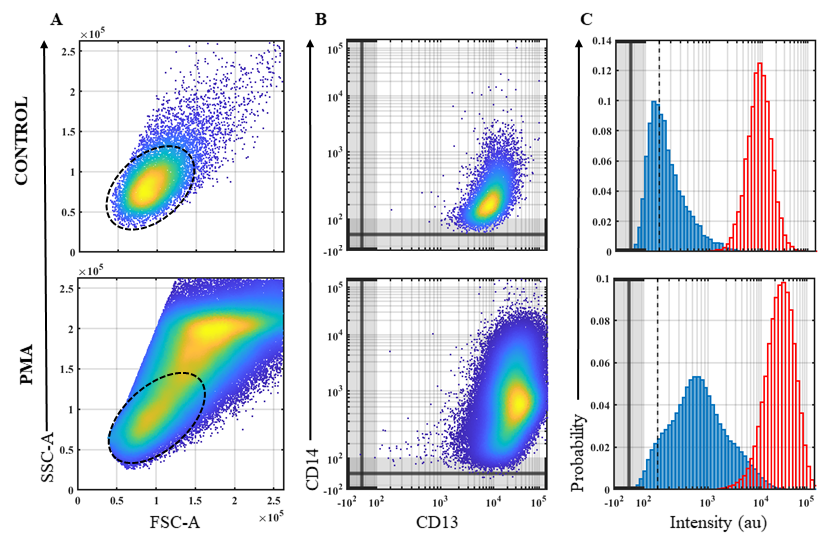

**Figure S5.** Phenotypical characterization of THP-1 cells prior to- (Control) and post- PMA-induced differentiation (PMA). PMA treatment renders cells larger in FSC-A and more granular in SSC-A (A), while expression of CD13 and CD14 markers increases compared to the control (B). The dashed oval gates highlight backgating of CD14(-) based on the vertical dashed gates at the histograms (C) of CD14 (bold blue) and CD13 (outlined red)

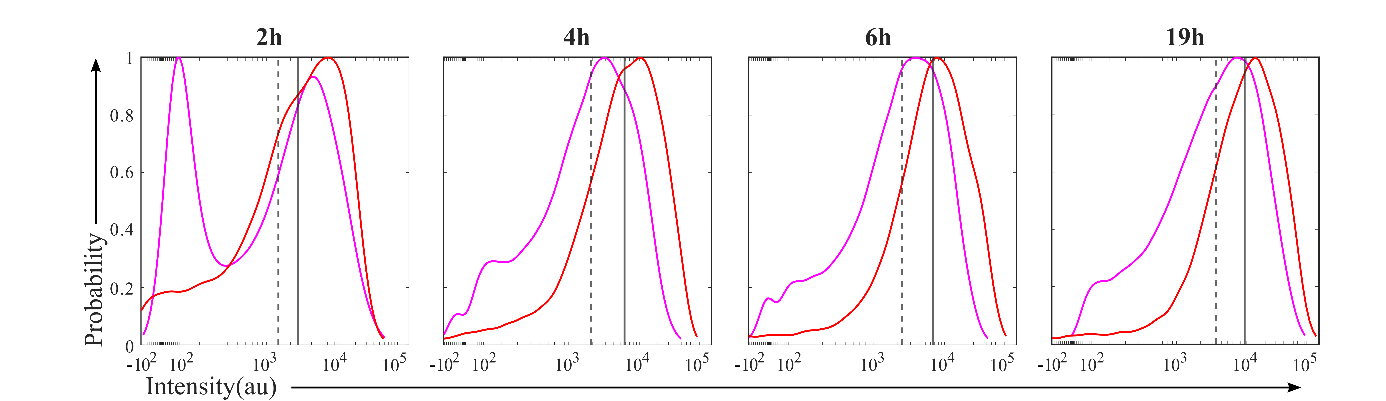

**Figure S6.** Histograms from single cell fluorescence analysis, by flow cytometry. Both monocytes (magenda) and monocyte-derived macrophages (red) were treated with NDs which exhibit increasing fluorescence signal (red) in the course of ND administration-time.The median ND fluorescence values are greater for the macrophages (gray bold) as opposed to the monocytes (gray dashed), at all time-points, indicating macrophages enhanced phagocytic capacity.
